# Evidence for rapid phenotypic and behavioral change in a recently established cavefish population

**DOI:** 10.1101/651406

**Authors:** Suzanne E. McGaugh, Sam Weaver, Erin N. Gilbertson, Brianna Garrett, Melissa L. Rudeen, Stephanie Grieb, Jennifer Roberts, Alexandra Donny, Peter Marchetto, Andrew G. Gluesenkamp

## Abstract

Substantial morphological and behavioral shifts often accompany rapid environmental change, yet, little is known about the early stages of cave colonization. Relative to surface streams, caves are extreme environments with perpetual darkness and low nutrient availability. The Mexican tetra (*Astyanax mexicanus*), has repeatedly colonized caves throughout Mexico, suggesting an ability to adapt to these conditions. Here, we survey for phenotypic and behavioral differences between a surface population and a cave population of *A. mexicanus* that has recently colonized Honey Creek Cave, Comal County, Texas, likely within the last century. We found that fish from Honey Creek Cave and fish from Honey Creek surface populations differ significantly in morphological traits including length, coloration, body condition, eye size, and dorsal fin placement. Cavefish also exhibit an increased number of superficial neuromasts relative to surface fish. Behaviorally, cavefish consume fewer worms when trials are performed in both lighted and darkened conditions. Cavefish are more aggressive than surface fish and exhibit fewer behaviors associated with stress. Further in contrast to surface fish, cavefish prefer the edges to the center of an arena and are qualitatively more likely to investigate a novel object placed in the tank. While cavefish and surface fish were wild-caught and developmental environment likely play a role in shaping these differences, our work demonstrates morphological and behavioral shifts for Texas cavefish and offers an exciting opportunity for future work to explore the genetic and environmental contributions to early cave colonization.

## Introduction

Colonization of new environments and other rapid changes to an organism’s environment offer unique opportunities to gain important insights into the role of genetics and plasticity in shaping phenotypes and behaviors (Atwell et al., 2012; Colautti and Lau, 2015; Gordon et al., 2009; Kinnison and Hairston Jr, 2007; Møller, 2010). For example, extensive evidence points to new environments eliciting evolutionary responses on contemporary timescales in response to environmental shifts (Dargent et al., 2019; Gordon et al., 2009; Hendry and Kinnison, 1999; Messer and Petrov, 2013; Reznick and Ghalambor, 2001; Whitehead et al., 2017). In addition, by reducing the strength of selection imposed by novel environments, plasticity can promote colonization of new environments (Ghalambor et al., 2007; Gimonneau et al., 2010; West-Eberhard, 2005), and this may be especially true for behavioral traits, which may be more environmentally labile than morphological traits (Baños-Villalba et al., 2017; Foster, 2013; Wcislo, 1989; West-Eberhard, 1989; Zuk et al., 2014).

One dramatic environmental comparison is between cave and surface environments. Caves are challenging environments due to perpetual darkness, absence of important environmental cues, low nutrient levels, and elevated levels of CO_2_ (Howarth, 1993; Poulson and White, 1969). The cave environment may also provide a refuge from predation and competition, seasonal and acute weather extremes, and UV radiation. Despite extreme differences in the selective landscape, cave-dwelling animals are numerous and caves are repeatedly colonized (Culver and Pipan, 2009; Howarth and Moldovan, 2018; Pipan and Culver, 2012). Cave organisms typically display a suite of characters including reduction of eyes and pigmentation (Culver and Pipan, 2009; Keene et al., 2015; Romero, 2009; Romero and Paulson, 2001), decreased metabolic rate (Hadley et al., 1981; Huppop, 1986; Niemiller and Soares, 2015), increased starvation tolerance (Hervant et al., 1999; Hervant et al., 2001; Huppop, 1986), and enhanced non-visual senses and associated structures (Bibliowicz et al., 2013; Hüppop, 1987; Protas et al., 2008; Yoshizawa et al., 2010). Many of these changes have evolved convergently in a diverse array of troglobites (Howarth, 1993; Juan et al., 2010; Niemiller and Soares, 2015; Pipan and Culver, 2012; Protas and Jeffery, 2012), yet, we do not understand the rate and sequence of these changes (and if they are comparable across taxonomic groups), nor the role of plasticity and evolved genetic changes in shaping the first cave-derived traits.

The Mexican tetra, *Astyanax mexicanus*, is a commonly used model organism for the study of vertebrate development, biomedical research, and adaptation to cave environments (Jeffery, 2001; Krishnan and Rohner, 2017; McGaugh et al., 2014; O’Quin and McGaugh, 2015). At least 29 populations of *A. mexicanus* have persisted in caves in the Sierra de El Abra and Sierra de Guatemala region of northeastern Mexico for hundreds of thousands of years and offer a natural laboratory for the study of evolution in cave environments (Gross, 2012; Herman et al., 2018; Mitchell et al., 1977). Individuals from cave populations display convergent phenotypes including reduction or loss of eyes and pigmentation, more posterior dorsal fin placement, shorter body length, and greater length-standardized body mass relative to surface fish (Protas et al., 2008). Differences in behavioral traits between cave and surface populations suggest a cavefish behavioral syndrome that includes lack of schooling (Kowalko et al., 2013), increased wall-following behavior (Sharma et al., 2009), reduced total sleep (Duboué et al., 2011; Jaggard et al., 2017; Jaggard et al., 2018), reduced stress (Chin et al., 2018), increased or decreased food consumption compared to surface fish (Tinaja and Pachón, respectively (Aspiras et al., 2015)), and reduced aggression (Burchards et al., 1985; Elipot et al., 2014; Elipot et al., 2013; Espinasa et al., 2005; Hinaux et al., 2015; Langecker et al., 1995; Rétaux and Elipot, 2013). Long-established cave populations of *A. mexicanus* are extensively studied (Gross, 2012; Herman et al., 2018; Ornelas-García et al., 2008; Ornelas-García and Pedraza-Lara, 2015), but to our knowledge no work has been conducted on recently established cave populations. Our study takes advantage of a recently-discovered, non-native cave population of *A. mexicanus* in Honey Creek Cave, Comal County, Texas which likely colonized the cave within the past century (Figure 1a).

**Figure 1.**
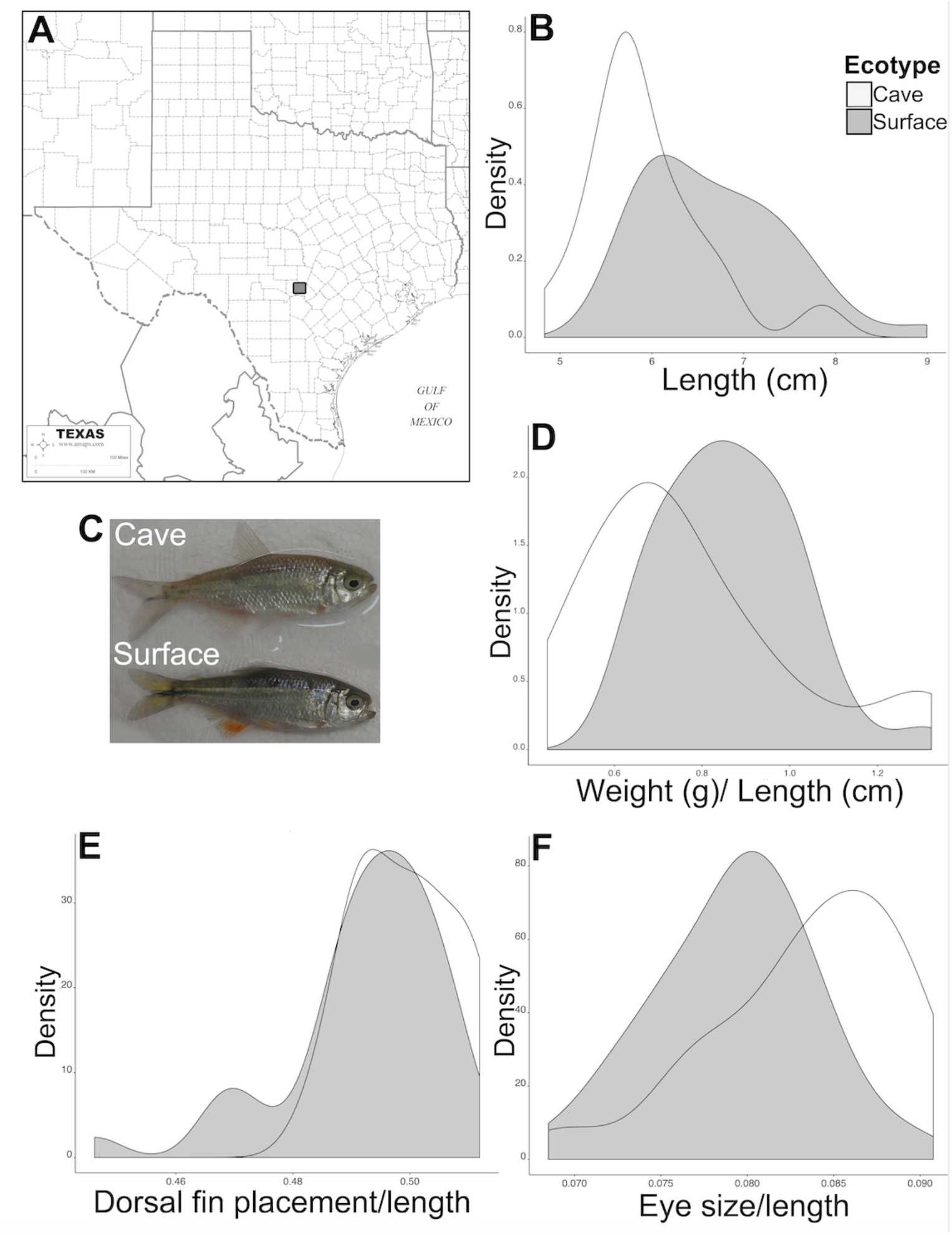
A) General location of Honey Creek Cave and surface sampling localities. B) Length differences between cave and surface (mean = 5.93 cm and 6.68 cm, respectively). C) Pictures of live cave and surface fish. D) Weight standardized by length of the fish (mean = 0.77 g/cm and 0.86 g/cm, respectively). E) Location of anterior insertion of the dorsal fin standardized by length of the fish (mean = 0.499 and 0.492, respectively). F) Eye diameter standardized by length of the fish (mean cave = 0.0834 and surface 0.0793).

Honey Creek Cave is the longest known cave in Texas (>30 km, (Veni, 1994)) and is part of the Edwards-Trinity aquifer system, which spans about 109,000 square kilometers of Texas (Barker and Ardis, 1996). With at least 91 cave species and subspecies (Bowles and Arsuffi, 1993; Culver et al., 2000), this aquifer system is one of the most biodiverse subterranean systems in the world (Longley, 1981). It is home to many endemic organisms, including the Comal blind salamander (*Eurycea tridentifera*), for which Honey Creek Cave is the type locality (Mitchell and Reddell, 1965). Although *Astyanax mexicanus* is native to the Rio Grande, Nueces, and Pecos rivers in South Texas (Mitchell et al., 1977), it was likely introduced to Central Texas in the early part of the last century as fish bait. The earliest record of *A. mexicanus* in the Guadalupe River Basin, which includes Honey Creek, was in 1953 (Constable et al., 2010), and the earliest observations of *A. mexicanus* in Honey Creek Cave were likely in the 1980s (Alan Cobb and Linda Palit, pers. comm), though biological investigations of the cave have been conducted since the early 1960s.

We assayed the Honey Creek Cave population for evidence of phenotypic and behavioral differences relative to a geographically proximate (< 1 km) Honey Creek surface population. We compared whether differences between Honey Creek Cave and surface populations are congruent with those observed between surface fish and long-established cave populations of *A. mexicanus* in the Sierra de El Abra and Sierra de Guatemala in Mexico. Our study lays a foundation for future work to explore the genetics and phenotypic plasticity underpinning the observed diverging traits in the early stages of cave colonization.

## Methods

### Honey Creek sampling and data collection

Individuals were collected from Honey Creek Cave and Honey Creek in the Guadalupe River Basin in Comal County, Texas from 21 - 25 May 2018 and 29 June - 3 July 2018 (Figure 1a, Figure S1). We sampled using dip nets (1/16 inch mesh) and collapsible prawn traps baited with canned cat food and/or sardines. All fish were collected in accord with UMN IACUC protocol #1705-34800A and were shipped to UMN via DeltaDash Cargo Services. Upon arrival, fish were transferred to 37.9 and 75.7L tanks and kept at a density of < 1 fish for every 6L of water on a 10:14 light cycle with lights on at 0800hrs. Fish were fed frozen and dried bloodworms or brine shrimp *ad libitum* 1-2x per day. For all measurements, behavior assays, and quantification, the researcher was blind to ecotype identity. Sex of fish was difficult to discern and assignments were inconsistent between researchers, so we did not include it as a covariate in any tests.

### Museum collections

We examined specimens from the collections at the University of Texas-Austin (Biodiversity Collections) and at Texas A&M University (Texas A&M Biodiversity Research and Teaching Collections). We photographed museum individuals sampled from the Guadalupe River Basin in the Texas counties of Comal (N = 3), Hays (N = 28), Kerr (N = 27), Kendall (N = 10), and Victoria (N = 7) and include all data in the supplementary materials (Table S1, Figure S1). To limit the effect of multiple sampling years and localities with very low sample size, we only evaluated 15 individuals from Hays County collected in 1976 and 20 individuals from Kerr County collected in 1979, as the other collections were smaller sample sizes from different years. Notably, these samples were taken only shortly before *A. mexicanus* was first observed in Honey Creek Cave in the 1980s, allowing us to directly compare the contemporary cave population to the likely ancestral surface population. Collections were also made from Honey Creek Cave in 2017 (N = 18) and compared to live fish collected in 2018 to examine consistency between years.

### Morphological analyses

Photographs were taken of each specimen using a Sony DSC-RX100 20.2 MP Digital Camera leveled on a tripod, with a color standard and ruler in each photo. ImageJ v1.46r (Schneider et al. 2012) was used to record the following measurements (in mm): eye diameter, standard length (distance from tip of snout to the posterior end of the last vertebra), and distance to dorsal fin (distance from the tip of the snout to the anterior insertion of the dorsal fin). Mass was taken with an AWS-100 balance (capacity: 100g, graduation 0.01g).

Measurements of coloration were taken from photos of the right side of each fish. Using a similar procedure as in (McGaugh, 2008), photos were opened in Photoshop 2015.5, the image was flattened, and the eyedropper tool was used to sample 31×31 pixels for red, green, blue (RGB), and hue, saturation, and lightness (HSL) values from the Color Picker window from four landmarks across the fish body. These landmarks included 1) the anterior insertion point of the dorsal fin on the body, 2) tail fin junction to the body with the landmark placed within the black stripe in the center of the fin and body, 3) the anterior insertion point of the anal fin on the body, and 4) center of body. To properly position the fourth landmark, we selected the rectangle tool and drew a rectangle between the tail fin and dorsal fin landmarks. The lower right corner of the rectangle was used as the center body landmark and the rectangle was deleted before taking color measurements. We avoided any water spots, glare, or other abnormality on the fish or on the color swatch and made notes of any abnormality. RGB and HSL values were also taken for a color standard in each photo. Photographs and weights of each live fish were taken on the same day, directly after behavior and feeding assays (see below). In total, we analyzed photographs and weights from 19 live cavefish and 40 live surface fish. We also analyzed photographs from 18 dead cavefish and 13 dead surface fish, as physiological color change can impact observed color for live fish.

Neuromasts were stained using a procedure similar to previously published methods (Jaggard et al., 2017; Yoshizawa et al., 2010). Fish were submerged in conditioned, aerated water with 0.05% DASPEI (2-4-dimethylamino-N-ethylpyridinium iodide; Sigma Aldrich) to label superficial and canal neuromasts. After staining, fish were anesthetized in an ice bath of conditioned water until immobilized. Ice-chilled water was required to immobilize surface fish, whereas cavefish required that water temperature be > 7°C to survive. Images were acquired using a Nikon TE2000 inverted fluorescence microscope with a filter set for detection of GFP (excitation 450–490 nm, 500 nm DM and 520/30 em filter) through a Nikon 1x, 0.04na x objective, using a Hamamatsu Flash 4 V2 camera with a 500 ms exposure and 10x magnification, running Nikon Elements v5.11 software. Throughout image acquisition, fish gills were wet with cool conditioned water via a bulb pipette. Fish were immediately returned to aerated water from their home tank after being photographed. Due to concerns for fish survival, only the right side of the body was imaged.

Superficial neuromast number, size, and size of the cranial third suborbital bone (SO-3) were counted using a custom macro written in Fiji v2.0.0-rc-69/1.52n (supplemental materials). The macro counted neuromasts by delineating light intensity and absolute size thresholds. The steps of the macro include: 1) taking a duplicate of the image, then smoothing out the duplicate using the median value with radius of 15 pixels, and subtracting the original image from the smoothed-out duplicate to remove noise, 2) drawing the region of interest, and 3) adjusting the threshold for size to be (2000,15600 microns^2^) and circularity of the neuromast to be (0.6-1) in order to be counted automatically. Size and circularity were empirically determined to include as many true positives as possible while excluding false positives. Each macro-processed image was visually inspected and corrected, if needed. To account for size differences between cave and surface fish, neuromast size (in pixels) and number were each divided by the area of the polygon (in pixels) of the cranial third suborbital bone as a standardization prior to statistical analysis. We tested 14 cavefish and 29 surface fish.

### Behavioral data collection and analysis

We quantified behavioral differences between Honey Creek Cave and Honey Creek surface populations and selected assays for which data existed for the long-established *A. mexicanus* populations in northeastern Mexico. These assays included: 1) vibration attraction behavior (VAB), which is a proxy for sensing moving food objects (Yoshizawa et al., 2010), 2) amount of food consumed under both light and dark conditions, 3) fish movement and spatial tank usage (i.e., a proxy for stress levels (Chin et al., 2018)), and 4) aggression in response to a mirror.

Room temperature was 20-21°C for all behavior assays, with no additional heat provided to tank water. Wyze Cams (v2) were used to record fish behavior in all trials across all lighting conditions. These cameras have the ability to provide clear, infrared recording and wireless connection. One camera per fish was used, and we controlled the cameras wirelessly while in a separate room from the behavior trials. All statistical analyses were performed in R v 1.1.463 (Team, 2013).

The VAB assays were conducted in the dark and were followed by the feeding assays in the light on the same day. About 5 weeks later, feeding in the dark was conducted. About two weeks after the dark feeding trials, the stress assays were conducted in both the dark and the light, and were immediately followed by the mirror-elicited aggression assays in the light. On trial days, the light cycle did not resume automatically at 0800 and the dark period was extended into the morning.

#### Vibration Attraction Behavior assays

Fish were fasted for 48hr prior to the trials in individual 2.8 L Aquaneering system tanks with Stress Coat®-treated (API) fresh tap water, as in their home tanks, and given aeration. After ∼24 hr a 20% water change was performed to prevent the buildup of ammonia.

For VAB trials, each fish was transferred and allowed to acclimate for 1 hour into a 22 cm circular arena with 10 cm depth of water, following (Yoshizawa et al., 2010). Three trials were conducted for each fish in the following order: 1) no rod, 2) rod with no vibration, and 3) rod with ∼35Hz vibration. Trials were each three minutes long. Between each trial, a researcher needed to enter the room to either place the rod or turn on the vibration, and fish were given an additional five minutes acclimation prior to data recording. The researcher left the room for the recorded trials, and trials were conducted with the lights off in the room and infrared recording. Six fish were tested per batch, and we conducted trials on three batches per day (a total of 18 fish tested per day). Trial arenas were rinsed with fresh tap water and refilled with conditioned water between batches of fish.

The level of vibration we targeted elicited the peak response from cavefish in (Yoshizawa et al., 2010). Vibration apparatuses were fashioned with Yootop 1 kΩ ½ W Trimming Variable Resistors Potentiometers, BestTong 14000 RPM 2 Wires Miniature Micro Vibrating Vibration Vibrator Motor DC 3V, and USB A Male Adapter Cables for power (Figure S2a). They were mounted on standard ring stands with clamp holders and a thin metal wire was used to transmit the vibration into the water (Figure S2b). The iPhone application Tuner T1 was used prior to each vibration trial to ensure the rod was vibrating at ∼35 Hz. The frequency was tuned initially in the lab by using a pair of aluminum foil contacts connected to a 9 V battery and a handheld multimeter (Model 173, Fluke Manufacturing) set to frequency mode.

Vibration attraction behavior was quantified from the video. Within the arena, we drew a 8.9 cm circle that was centered on the location of the vibrating rod. An approach was defined as when a fish changed direction to swim towards the wire and reached within the radius of the inner circle to the wire. We recorded the number of approaches manually, and recorded the time within the inner circle, transitions in and out of the circle, total distance traveled, and velocity with Ethovision XT 14 (Noldus). In total, we tested 18 cavefish and 39 surface fish for VAB. For four cavefish and five surface fish, the metal rod was not placed perfectly over the center of the inner circle. We were able to quantify the number of approaches to the rod for these trials, but did not include them in the automated Ethovision analysis, resulting in a total of 14 cavefish and 34 surface fish for which the Ethovision-measured parameters were included in statistical analyses.

#### Feeding assays

Directly after the VAB trials, fish were given at least 1hr to acclimate back into their original 2.8L Aquaneering fasting tanks, with opaque separators between tanks. If repositioning tanks was needed for better observation, fish were given another 30 mins of acclimation. Fish were given an additional 10min of acclimation time if observers were added to the room. Frozen bloodworms were weighed to the nearest 0.01g. Researchers used tweezers to feed individual bloodworms to individual fish for 10 min, feeding an additional blood worm when one was consumed. We recorded the weight of the remaining bloodworms by reweighing the weighboat to the nearest 0.01g, the time to first feeding (latency), and counted how many bloodworms were consumed over a 10 minute period. We found that evaporation consistently led to unreliable weights of the blood worms, thus, we analyzed data for number of bloodworms consumed. We tested 19 cavefish and 40 surface fish.

Since the first feeding trial was conducted under conditions which might have inhibited a feeding response from cavefish (e.g. researchers were present for the first feeding trial, and it was conducted in the light), we conducted a second feeding trial several weeks later in the dark without researchers present. The same fish were used as before, but since fish were returned to their communal tanks between trials, we did not match fish identity between the first and second trials. Trials were conducted approximately at the normal “lights on” time of 0800. As Mexican cavefish are known to be more resilient to starvation (Aspiras et al., 2015), fish were fasted for approximately 120 hrs before this trial (three days longer than for the first feeding trial) to ensure we fasted the cavefish long enough to elicit a feeding response, and we recorded how many bloodworms were consumed over a 10 minute period. Each fish was supplied with 50 total bloodworms, and then the researcher left the room. Bloodworms remaining in the tank were counted after 10 minutes. Fish were weighed after the trial to account for fish size differences in the analysis. Over the course of three weeks, we tested 17 cavefish and 17 surface fish.

#### Stress assays

Cavefish experience a reduced predation risk compared to surface fish, thus, cavefish may demonstrate decreased behavioral responses to stressful stimuli (Chin et al., 2018). Indicators of stress rely on reduced exploratory behavior of a new environment, which includes: shorter distances traveled, longer durations of time spent in the bottom half of the tank, slower velocity, and longer periods of “freezing” immobility (Chin et al., 2018). Thus, we measured four stress behaviors: total distance traveled, velocity, duration of time spent in the bottom tank half, and duration of time spent immobile. Immobility state threshold was set as less or equal to 10.00% change in the complete area of the subject. We tested 15 cavefish and 37 surface fish.

Fish were allowed to acclimate in 18.9L tanks at room temperature (20-21°C) for 1 hr without aeration. After 1hr of acclimation, we conducted 5 min of recordings and analysis in the dark, and 5 min in the light, waiting for a full minute after lights were turned on to begin quantification of behaviors. To enhance the ability of Ethovision to recognize the fish, we used the differencing function under advanced detection settings to compare the video to a reference image without the subject. Acquisition resulted in < 9.2% ‘subject not found’ data for each video (median = 1.1%) in the dark and < 32.6% in the light (median = 5.25%), and we interpolated missing data using the Track Editor function. Incorrect subject tracking and interpolation data were also manually corrected using the Track Editor function so that the final measures contained no missing data.

#### Aggression assays

Directly after the stress assay, we conducted mirror-elicited aggression assays. A researcher entered the room and placed a mirror in the tank, and this mirror covered the entire short side of the aquarium except for a few cm at the top of the tank. Proportion of time spent within 15 cm of the mirror was quantified with Ethovision over the course of an hour-long trial. Similar methods for video acquisition and interpolation were used as in the stress assay except the differencing function and reference videos were not used for all videos. We tested 14 cavefish and 35 surface fish for aggression because we excluded three trials with > 20% missing data prior to interpolation (max in retained trials = 19.7%, median = 10.9 %).

## Results

### Morphological analysis

For length, weight, eye size, and coloration of live fish, 19 cavefish and 40 surface fish were included in the analyses presented below.

#### Cavefish are shorter and weigh less than surface fish

Honey Creek Cave fish were significantly shorter than Honey Creek surface fish (mean = 5.93 and 6.68 cm, respectively; Wilcoxon W = 145.5, p = 0.0004, Figure 1b). This is similar to that documented in (Protas et al., 2008), though, length distributions of wild collections can vary substantially when not age-matched. Examples of collected fish are given in Figure 1c. Honey Creek Cave fish weigh less per unit length than surface fish (mean = 0.77 g/cm and 0.86 g/cm, respectively; Wilcoxon W = 220, p = 0.0224, Figure 1d). This is not consistent with previous findings in Mexican cavefish, which typically weigh more than their surface counterparts (Aspiras et al., 2015; Protas et al., 2008; Riddle et al., 2018). Measurements were taken within two months of capture from the field, thus, these body conditions could potentially reflect lower resource availability in the caves relative to the surface populations.

#### Dorsal fin placement is more posterior in Honey Creek Cave fish

Previous work documented that the anterior insertion of the dorsal fin, when standardized for the length of the fish, is more posterior in cave populations of *A. mexicanus* than in surface populations (Protas et al., 2008). We standardized dorsal fin placement by dividing by standard length of each specimen, as done previously (Protas et al., 2008). We found that the insertion of the dorsal fin was more posterior in cave individuals than in surface individuals, though the effect size was not large and not statistically significant with a nonparametric test (mean = 0.499 and 0.492, respectively; Wilcoxon W = 451, p = 0.087; Welch Two Sample t-test t = 2.28, df = 50.887, p = 0.027) (Figure 1e).

#### Eye diameter is larger in Honey Creek Cave fish

Comparisons of length-standardized eye size revealed significant differences between cave and surface individuals. Individuals from Honey Creek Cave exhibited larger standardized eye size than surface individuals (mean = 0.0834 and 0.0793, respectively; Wilcoxon W = 520, p = 0.0031, Figure 1f). Notably, the difference in eye size between ecotypes is not present when the data are analyzed with a linear model with eye size as the response variable, length as a covariate, and ecotype as the factor. We suspect this is because the distributions of fish length are substantially different for ecotypes, thus, standardizing eye size for each fish by their length is more appropriate.

#### Honey Creek Cave fish exhibit higher saturation coloration than surface fish

Mexican cavefish are albinistic or exhibit reduced number and size of melanophores (Gross et al., 2009; Protas et al., 2006; Stahl and Gross, 2015). For four landmarks on the fish, we quantified HSL (Hue, Saturation, Lightness) and RGB (Red, Green, Blue) values from photographs of live fish (McGaugh, 2008; Sacchi et al., 2013). We analyzed all color data for all landmarks using a Principal Components Analysis (PCA) and determined that cave and surface fish were separated mainly by PC2 (Figure 2a). Next, we analyzed each metric for all four landmarks using PCAs and determined that saturation was likely driving the majority of the signal from the PCA of all color components (Figure 2b). Higher saturation values represent colors with less gray components, which could be interpreted as less dark pigmentation.

**Figure 2.**
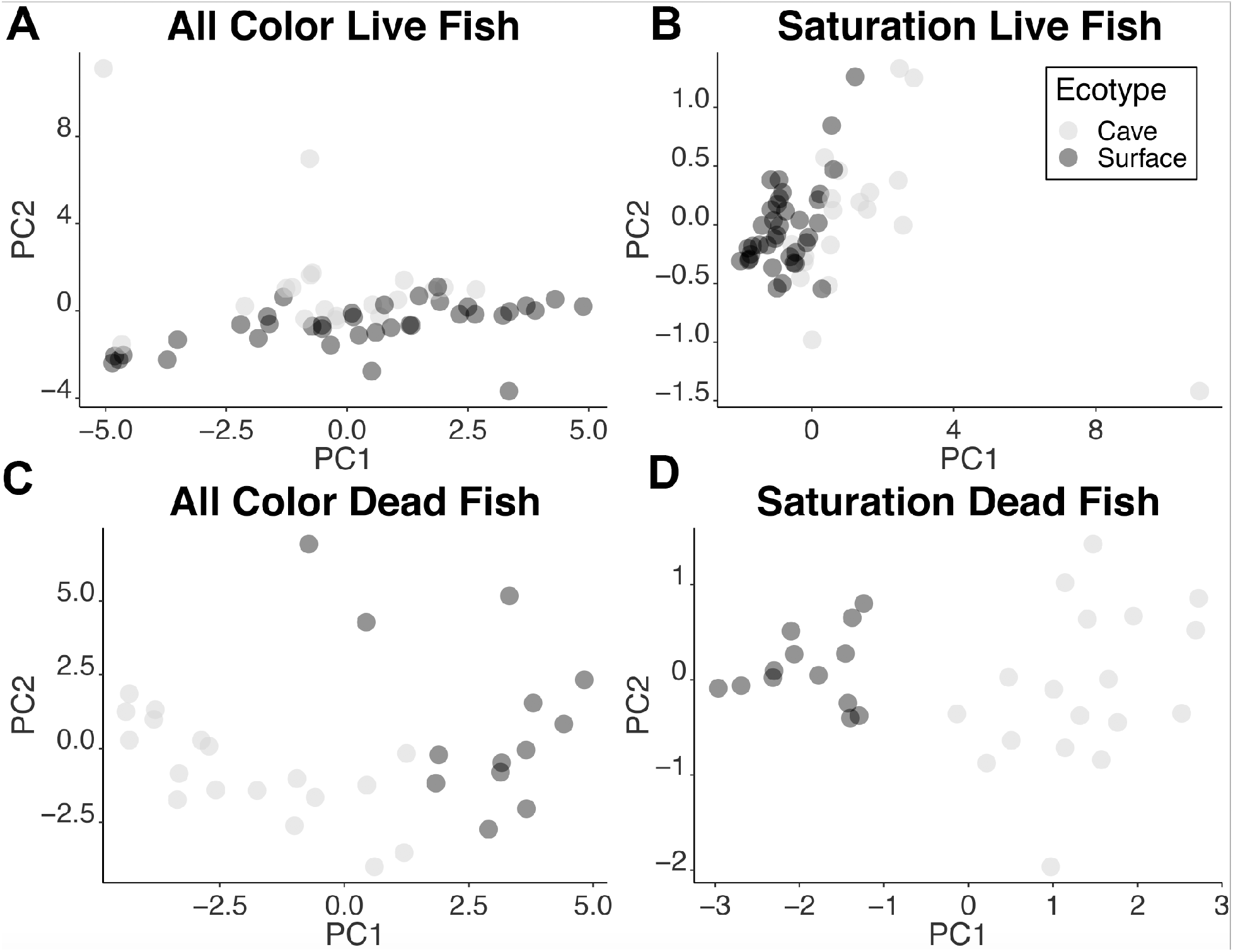
A) PCA of all color components of live fish (N =19 cave, 37 surface) B) Saturation color component of live fish (N =19 cave, 37 surface), C) PCA of all color components of dead fish (N =18 cave, 13 surface), D) Saturation of dead fish (N =18 cave, 13 surface).

Rapid physiological color change occurs in fish and can potentially interfere with color analyses of live fish (Sköld et al., 2013), overall, the dead fish (18 cave, 13 surface) corroborate color shifts between the two ecotypes. Separation with PCAs based on all color components and saturation alone was evident for dead cave and surface fish (Figure 2c, 2d).

Cavefish consistently exhibit higher saturation values than surface fish (p < 0.001, for all four landmarks) for both live and dead fish (Table S2, S3). No metrics aside from saturation are significant across all four landmarks. Dead cavefish exhibited significantly greater lightness values for anal and tail landmarks than dead surface fish, but the reverse is true for the middle body landmark (Table S3). Overall, our analysis suggests that Honey Creek Cave fish exhibit coloration shifts toward less dark pigmentation, concordant with Mexican cavefish (Culver and Pipan, 2016; Gross et al., 2009; Howarth and Moldovan, 2018; Kronforst et al., 2012; Pipan and Culver, 2012; Protas et al., 2007; Protas et al., 2006).

#### Museum collections confirm dorsal fin placement shift

For dorsal fin placement, preserved surface fish collected in the 1970s from Hays County and Kerr County exhibited more anteriorly-set dorsal fins than Honey Creek Cave fish collected in 2018, but also more anteriorly-set dorsal fins than Honey Creek surface fish collected in 2018 from Comal County (mean Hayes = 0.437, Kerr = 0.444, HCC_2018_ = 0.499, HC surface_2018_ = 0.491, p < 0.0001 for all Wilcoxon rank sum tests). Honey Creek Cave fish collected in 2018 exhibited more posterior-set dorsal fins compared to those collected in 2017 (HCC_2017_ = 0.474, HCC_2018_ = 0.499, p < 0.001). Both 2018 and 2017 Honey Creek Cave fish exhibited more posterior-set dorsal fins than both museum surface groups from 1976 and 1979.

Eye size per unit length (HCC_2017_ = 0.087) was not significantly different between 2018 and 2017 collections of Honey Creek Cave fish. Hays Co. surface fish exhibited smaller eyes per unit body length than 2018 Honey Creek Cave fish and Honey Creek surface fish (mean Hayes = 0.069, HCC_2018_ = 0.084, HC surface_2018_ = 0.078, p < 0.0001 for all Wilcoxon rank sum tests). In contrast, Kerr Co. surface fish exhibited nearly identical eye size per unit body length (Kerr = 0.086) compared to Honey Creek Cave fish (p = 0.184) and larger eyes per unit body length compared to Honey Creek surface fish (p < 0.001). This suggests that the eye size shifts observed for Honey Creek Cave fish are well within natural surface variation, may not be in response to the cave environment, and all features should be interpreted with year-to-year variation in mind.

#### Cavefish have more suborbital superficial neuromasts

We observed that Honey Creek Cave fish (N = 14) possessed about 1.3-fold the number of suborbital superficial neuromasts than surface fish (N = 29), after dividing the neuromast count by the size of the cranial third suborbital bone (Figure 3a, Wilcoxon W = 330, p = 0.0007). In absolute numbers, cavefish have about 1.05-fold the number of neuromasts (mean: cave =136, surface =131, median: cave = 135, surface = 126), but the area of the suborbital of cavefish is 77% the size of the surface fish (cave = 532312.3 pixels, surface = 684035.0 pixels). Thus after accounting for the size of the focal area, cavefish exhibit a substantial and significant increase in neuromasts relative to surface fish.

**Figure 3.**
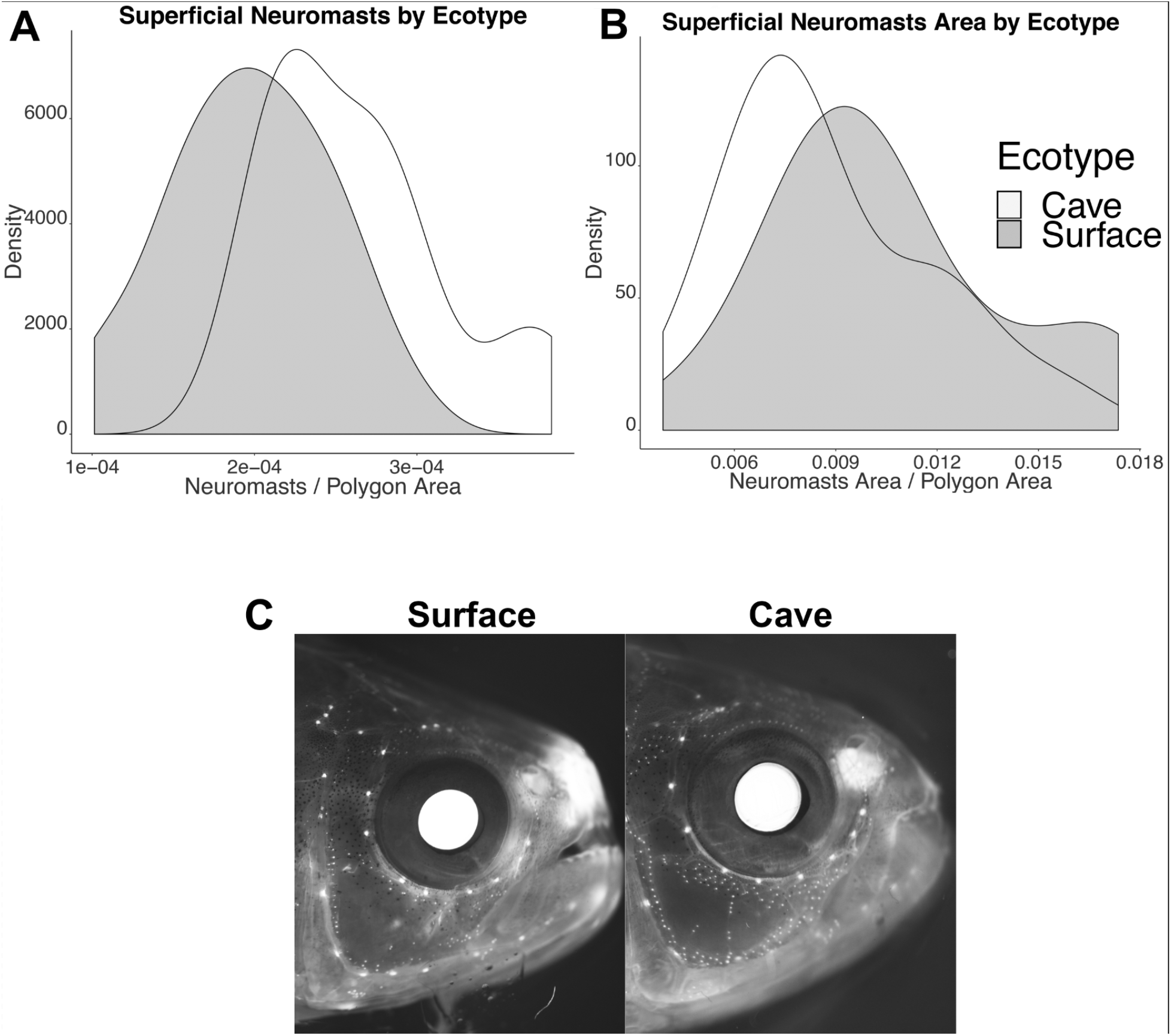
A) Number of superficial neuromasts standardized by the size of the suborbital bone. B) Mean area of superficial neuromasts standardized by the size of the suborbital bone. C) Images were picked from the ends of the distribution for number of neuromasts. Original images were adjusted image-wide for brightness and contrast only. N = 14 cavefish, N = 23 surface fish.

We observed a qualitative trend of smaller neuromasts in cavefish relative to surface fish when comparing mean or median neuromast size per fish (divided by the size of the cranial third suborbital bone), but this was not significant (Figure 3b, mean: Wilcoxon W = 124, p = 0.257; median: W = 113, p = 0.138). Our observations (Figure 3c) are concordant with observations in Mexican cavefish which have more numerous neuromasts than surface fish in the area delineated by cranial third suborbital bone. However, Mexican cavefish also exhibit larger neuromasts than Mexican surface fish.

### Behavioral analysis

#### Cavefish and surface fish do not display VAB. Cavefish prefer the edges of the arena and do not avoid novel objects

To assess VAB, we analyzed the number of approaches to a rod and the proportion of the trial the fish spent in the inner circle of the arena for 18 cavefish and 39 surface fish. For the first measure of VAB, the number of approaches was qualitatively larger for cavefish in trials with the plastic rod without vibration (Trial 2: Cave mean = 5, Surface mean = 4.2; Cave median = 5, Surface median = 3, Figure 4a) and in trials with the rod vibrating at 35 Hz (Trial 3: Cave mean = 4.7, Surface mean = 4; Cave median = 4.5, Surface median = 4, Figure 4b). However, the larger number of approaches for cavefish was not statistically significant (Trial 2: Wilcoxon W = 391.5, p = 0.489; Trial 3: Wilcoxon W = 373.5, p = 0.704).

**Figure 4.**
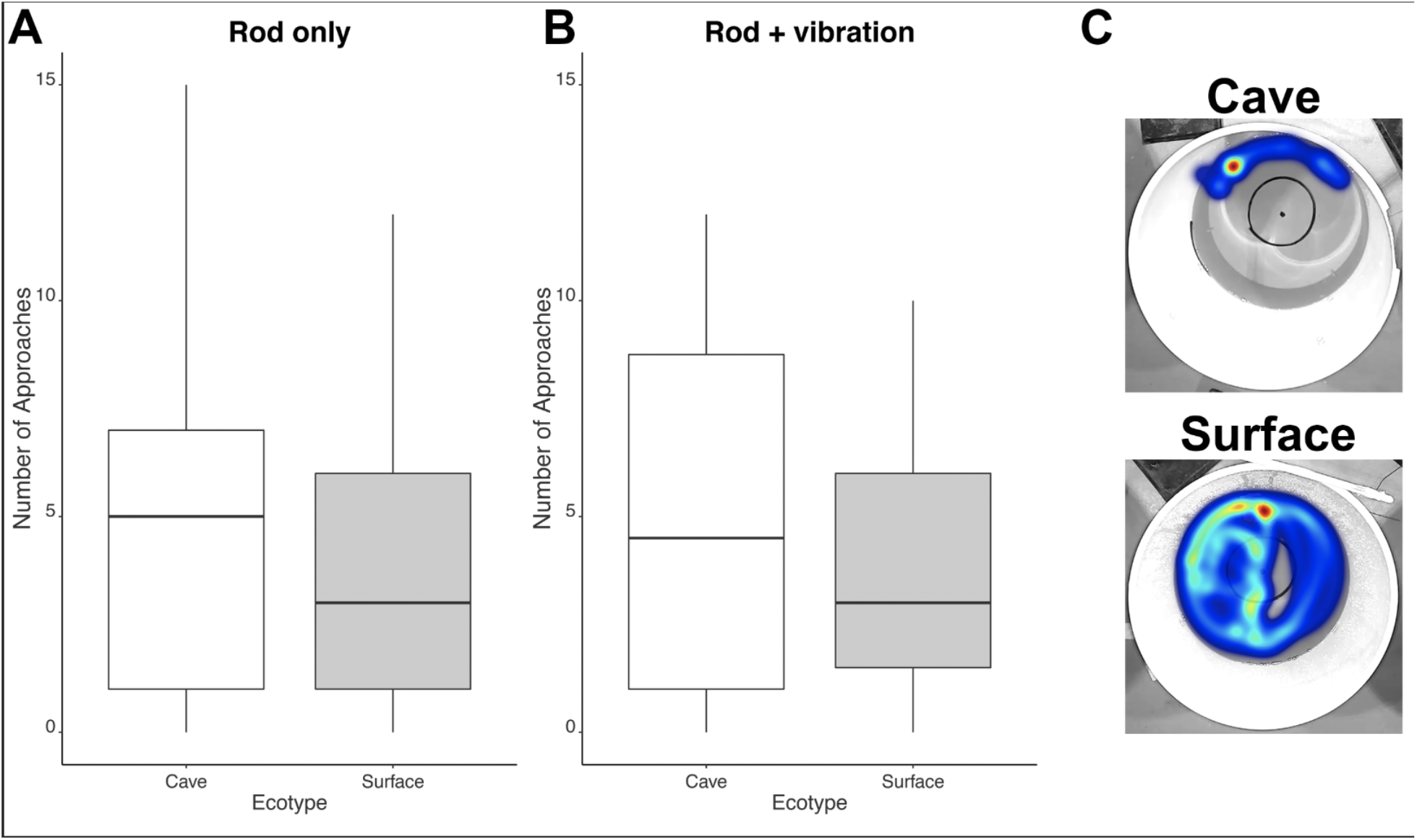
A) Number of approaches to a non-vibrating rod across a three minute trial period. B) Number of approaches to a rod vibrating at 35Hz across a three minute trial period. Both trials were conducted in the dark with the vibrating rod trial occurring immediately after the non-vibrating rod trial (N = 18 cave, 39 surface for both trials). C) Heatmaps from a trial with no rod (Trial 1) from cave and surface fish. Images were picked from the opposite ends of the distribution for number of crossings of the inner circle.

We also observed no difference between the number of approaches between Trial 2 (rod only) and Trial 3 (rod + vibration) for either ecotype (paired one-tailed Wilcox tests, p > 0.57, both cases, Figure 4). Thus, fish did not increase the number of approaches once vibration was added, regardless of ecotype (Table S4).

For a second measure of VAB, we recorded the proportion of the trial the fish was in the inner circle for 14 cavefish and 34 surface fish. Cavefish spent considerably less of their time in the inner circle compared to surface fish for all three trials. This difference was statistically significant for Trial 1 (no rod: Wilcoxon W = 83.5, p-value < 0.001) and Trial 3 (rod with vibration: Wilcoxon W = 143.5, p-value = 0.033). The lack of statistical significance in Trial 2 was likely due to a lack of statistical power, as cavefish were located in the center of the arena only about 60% as often as surface fish. We interpret these results as stronger wall-following behavior in cavefish relative to surface fish, which has been documented previously (Sharma et al., 2009). This is further supported by our observation that cavefish exhibit significantly fewer transitions in and out of the center circle than surface fish for Trial 1 (no rod: Wilcoxon W = 138.5, p-value = 0.024; Cave median = 6.5, Surface median = 11; Figure 4c) and qualitatively fewer transitions for Trial 3 (rod with vibration: Wilcoxon W = 181.5, p-value = 0.203; Cave median = 4.5, Surface median = 7.5).

This difference between cave and surface fish behavior appears to be due mainly to a decrease in surface fish transitions and a decrease in the amount of time surface fish spent in the center of the arena once the rod was added, rather than a change in cavefish behavior (Paired one-tailed Wilcoxon tests, Table S4). Taken together, these results suggest that cavefish prefer to be located near the margins of the arena (similar to (Patton et al., 2010; Sharma et al., 2009)), but are qualitatively less likely than surface fish to avoid novel objects (similar to (Yoshizawa et al., 2010), which observed shorter latency to approach the non-vibrating rod).

#### Cavefish eat less than surface fish

Populations of Mexican cavefish vary in their feeding behavior, as they have been documented to eat significantly more (e.g. Tinaja) and significantly less (e.g. Pachón) compared to surface fish populations (Aspiras et al., 2015). We conducted two rounds of feeding trials with Honey Creek cave and surface fish. First, after 48h of fasting, we placed a blood worm with individually-housed fish (19 cavefish, 40 surface fish) and waited for the fish to consume the worm before adding another. This was performed in a lighted room. We found cavefish ate significantly fewer worms than surface fish over a 10-minute period after dividing the number of worms eaten by mass of the specific fish (mean cavefish = 4.63 total worms [1.21 worms/g of fish], surface fish = 26.81 total worms [4.70 worms/g of fish]; Wilcoxon W = 122, p < 0.0001, Figure 5a).

**Figure 5.**
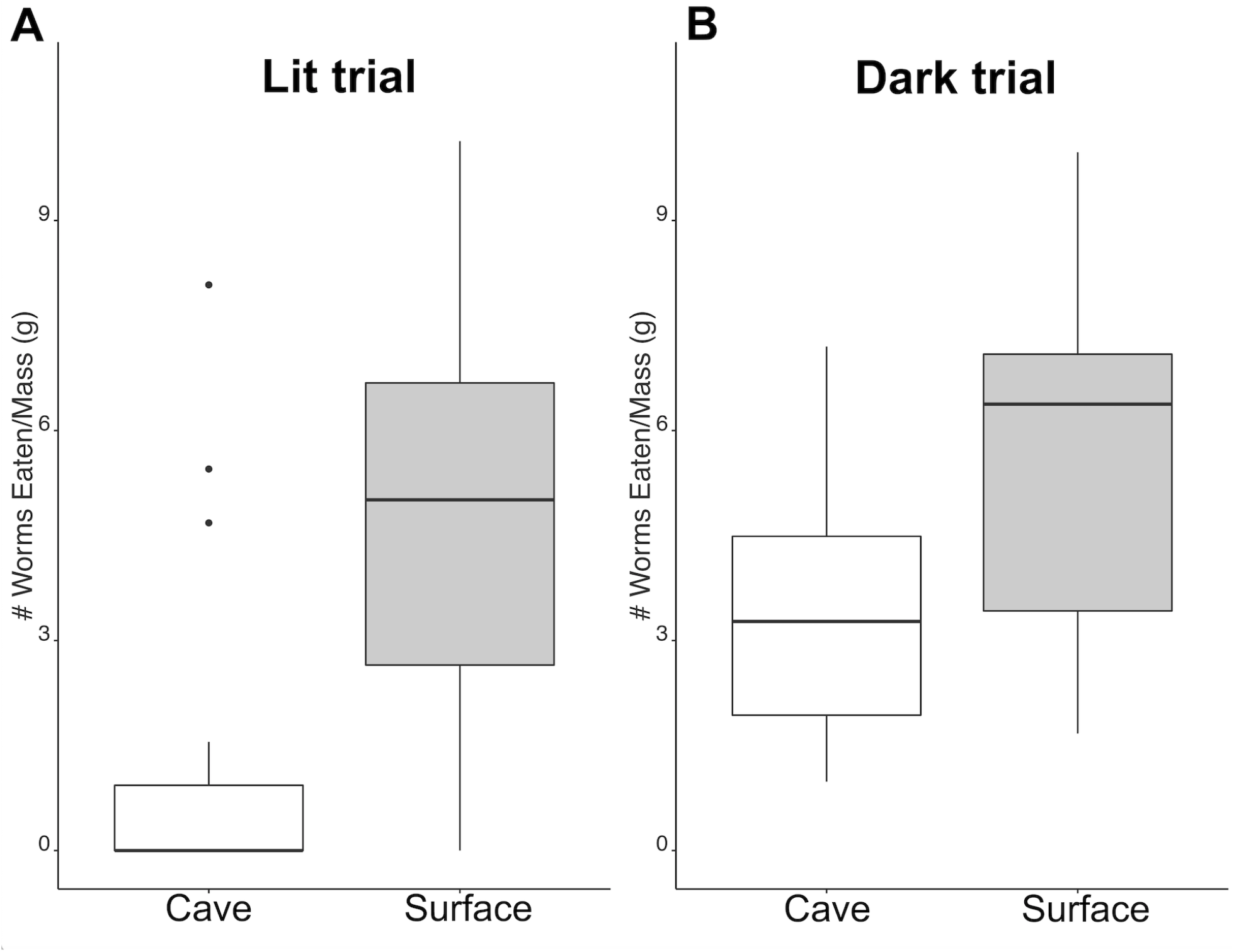
A) Number of worms consumed over a 10 minute period after a 48h fasting period conducted in a lighted room with researcher present (N = 19 cave, 37 surface). B) Number of worms consumed over a 10 minute period after a 120h fasting period conducted in the dark with no researcher present (N = 17 cave, N = 17 surface).

Second, we repeated feeding trials with a longer fasting time prior to the trial (120h) and conducted the trials in the dark without a researcher present using 17 cavefish and 17 surface fish. As cavefish may be less likely than surface fish to eat when researchers were present and could be less affected by fasting conditions than surface fish (Aspiras et al., 2015), our goal with the second round of feeding trials was to reduce the impact of these variables. Cavefish ate significantly fewer worms than surface fish over a 10 minute period after dividing the number of worms eaten by the mass of the specific fish (mean cavefish = 14.94 worms, surface fish = 31.12 worms; Wilcoxon W = 72, p < 0.012, Figure 5b). Thus, it appears lower feeding rates are a consistent trait of Honey Creek Cave fish relative to surface fish.

#### Surface fish exhibit more stress than cavefish

We examined the average distance, average velocity, proportion of time in the lower half of the tank, and proportion of time spent immobile for 15 cavefish and 37 surface fish for 5 min under dark conditions and 5 min under lighted conditions. We found no difference in any of these metrics during the dark trial (Wilcoxon rank sum tests, distance: W= 250, p = 0.589; velocity: W = 253, p = 0.631; lower ½ of tank: W = 213, p = 0.198; immobile: W = 274, p = 0.952; results consistent if tested with parametric t-test) or light trial (Wilcoxon rank sum tests, distance: W= 317, p = 0.435; velocity: W = 340, p = 0.213; lower ½ of tank: W = 299, p = 0.675; immobile: W = 225.5, p = 0.298; results consistent if tested with parametric t-test).

However when the dark and light trials were analyzed by a paired Wilcoxon test, some interesting differences emerged. Notably, in the light, surface fish significantly slowed their velocity (V = 501, p = 0.023) and reduced their distance traveled (V = 509, p = 0.0166) by about 85%. In contrast, cavefish did not change their velocity or distance traveled once the lights were turned on (V = 56, p = 0.847; V = 66, p = 0.762, respectively). Both cave and surface fish spent significantly more time at the bottom of the tank in the light trials than in the dark trials (Table S5; p < 0.0001 in both cases). These results suggest that cavefish exhibit fewer stress behaviors than surface fish.

#### Cavefish are more aggressive than surface fish

In the long-established Mexican populations, surface fish are more aggressive that cavefish (Elipot et al., 2013; Rétaux and Elipot, 2013). In contrast, Honey Creek Cave fish spent 85.1% of their time in the 15cm zone closest to the mirror, whereas Honey Creek surface fish spent 64.8% of their in the same zone (W = 349, p = 0.021; 14 cavefish and 35 surface fish). Fish generally adhered close to the mirror and appeared to pace vertically. We observed very few ramming motions which would be typical of two fish interacting in a tank.

## Discussion

In this study, we found evidence for shifts in morphological and behavioral traits in a recently-established cave population of *Astyanax mexicanus* relative to a geographically proximate surface population. Our study is the first to examine phenotypes of a recently-established *Astyanax mexicanus* cave population. Repeated evolution can tell us much about the evolutionary process, but complicating factors such as gene flow between populations can limit the strength of inference about the deterministic and predictable nature of evolution and natural selection (Rosenblum et al., 2014; Stern, 2013). One advantage of studying the Honey Creek Cave population is that it is highly unlikely that gene flow is transporting cave-adapted alleles from Mexican caves to Texas caves. Thus, if the morphological and behavioral differences described in this work are maintained after breeding in the laboratory in a common garden experiment, this population will provide a valuable, truly novel origin of trogolomorphy in which to study repeated evolution and an opportunity to explore selection and plasticity in the early stages of cave colonization.

Although *Astyanax mexicanus* is native to the Rio Grande, Nueces, and Pecos rivers (Mitchell et al., 1977), it was likely introduced to Central Texas in the early part of the last century. The earliest records for Central Texas come from fish hatcheries in Kerrville and San Marcos, and *A. mexicanus* may have been inadvertently collected along with gamefish stock sourced from within the native range of *A. mexicanus*. Brown (1953) reported deliberate introductions in San Pedro Springs (San Antonio River system) and the San Marcos River in 1908 and 1928-1930, respectively. Later, range expansion of this species into other local rivers was facilitated by entrepreneurs who collected *A. mexicanus* from the Rio Grande for sale as bait, beginning in the 1950s (Chester Critchfield, pers comm). Published in 1952, routine surveys reported no *Astyanax* present in Comal Springs and the Comal River, a tributary of the Guadalupe River (Ball et al., 1952), and 1953 was the earliest museum record of *A. mexicanus* in the Guadalupe River (VertNet, (Constable et al., 2010)). This species is currently abundant at those sites, thus, it is likely that it may have only recently invaded or was introduced into the Guadalupe River Basin.

While *A. mexicanus* may have invaded Honey Creek Cave soon after its range expanded into the Guadalupe River Basin, the earliest observations of *A. mexicanus* in Honey Creek Cave were in the 1980s (Alan Cobb and Linda Palit, pers. comm), despite biological surveys of Honey Creek Cave reaching back to the early 1960s. James Reddell, who collected and described the Comal blind salamander in the 1960s, did not recall observing *A. mexicanus* in the cave during that time (Reddell, pers comm). Though speculative, it is possible that extreme flooding of the Guadalupe River Basin from Tropical Storm Amelia (1978) catalyzed *A. mexicanus* colonization of Honey Creek Cave. Since the original observations in the 1980s, casual observations have documented the persistence of *A. mexicanus* in Honey Creek Cave. While historical records suggest a compelling case for a recent invasion, without genetic data we cannot fully rule out a Pleistocene refugia origin for this population.

Some of the observed shifts in Honey Creek Cave fish parallel those found in long-established Mexican cave populations (Table 1). First, Honey Creek Cave individuals have slightly more posterior dorsal fin location than their surface-dwelling counterparts. The dorsal fin serves as an important stabilizer and force generator for fish locomotion (Drucker and Lauder, 2001, 2005; Liao, 2007; Standen and Lauder, 2005, 2007); thus, the shift towards a more posterior dorsal fin may be in response to the slower flowing water or less predation pressure in caves and is consistent with long-established Mexican cave populations (Protas et al., 2008). Differences between individuals from Honey Creek Cave and surface fish was not consistent in Honey Creek Cave samples from 2017, though, both 2017 and 2018 Honey Creek Cave fish exhibited more posterior-set dorsal fins than surface fish collected in 1976 and 1979.

**Table 1.** Traits for *Astyanax mexicanus* inhabiting Sierra de El Abra and Guatemala regions of Northeastern Mexico and comparisons to Texas Honey Creek Cave and surface fish. Traits in bold are concordant between the two comparisons.

| **Trait** | **Mexican cavefish vs. surface** | **Texas cavefish vs. surface** |
| --- | --- | --- |
| **Length** | **Surface > Cave (Protas et al., 2008)** | **Surface > Cave** |
| **Coloration** | **Surface > Cave (Protas et al., 2008)** | **Surface > Cave** |
| **Feeding** | **Cave-specific (Aspiras et al., 2015)** | **Surface > Cave** |
| **Stress** | **Surface > Cave (Chin et al., 2018)** | **Surface > Cave** |
| **Wall-following** | **Surface < Cave (Sharma et al., 2009)** | **Surface < Cave** |
| **Neuromast number** | **Surface < Cave (Yoshizawa et al., 2010)** | **Surface < Cave** |
| **Dorsal Fin** | **Surface < Cave (Protas et al., 2008)** | **Surface < Cave** |
| Mass/Length | Surface < Cave (Protas et al., 2008) | Surface > Cave |
| Eye diameter/Length | Surface > Cave (Protas et al., 2008) | Surface < Cave |
| Aggression | Surface > Cave (Rétaux and Elipot, 2013) | Surface < Cave |
| VAB | Surface < Cave (Yoshizawa et al., 2010) | Surface = Cave |
| Neuromast size | Surface < Cave (Yoshizawa et al., 2010) | Surface = Cave |

Second, individuals from Honey Creek Cave exhibit lighter coloration (likely due to reduced melanization) than surface fish in comparisons of both live and dead fish, suggesting that this observation is not driven simply by physiological color response. Reduced pigmentation is one of the most commonly observed troglomorphic traits across cave-dwelling taxa (Culver and Pipan, 2016; Gross et al., 2009; Howarth and Moldovan, 2018; Kronforst et al., 2012; Pipan and Culver, 2012; Protas et al., 2007; Protas et al., 2006). Lighter coloration in the cave environment may be shaped predominantly by drift (Borowsky, 2015). Yet, other work suggests that coloration can be pleiotropically related to basic physiological processes (Ducrest et al., 2008; Roulin and Ducrest, 2011) and may be advantageous in the cave environment (Bilandžija et al., 2018; Bilandžija et al., 2013). It is intriguing that reduced dark pigmentation is among the earliest phenotypic changes observed in the recently established Honey Creek Cave population.

As in Mexican cavefish (Gertychowa, 1970; Patton et al., 2010; Sharma et al., 2009), Honey Creek Cave fish exhibit a stronger preference to the outer edge of the arena than surface fish as shown by our VAB trials. Despite spending less time in the center of the arena than surface fish, cavefish approached the rod at equal or higher rates, in part, because surface fish reduced their time in the center of the arena after addition of a novel object. These two observations suggest that Honey Creek Cave fish may exhibit more exploratory behavior than surface fish (Sharma et al., 2009), which is similar to observations in colonizers of other species (Candler and Bernal, 2014). Increased exploratory behavior in Honey Creek Cave fish also seems to be supported by cavefishes’ tendency to adhere more to the mirror during aggression trials. This increased exploratory behavior may be influenced by reduced predation in the cave environment, and reduced stress-related behaviors in Honey Creek Cave fish in lighted trials support this notion (Chin et al., 2018).

Honey Creek Cave fish eat substantially less than surface fish under multiple conditions. This is similar to comparisons of Pachón cavefish to Mexican surface fish (Aspiras et al., 2015). Differences in food consumption and reduced body condition (mass per unit length) suggest that metabolic shifts in Honey Creek Cave fish should be investigated in future work, as low and unpredictable food availability is a major selection pressure in many caves (Huppop, 2000; Niemiller and Soares, 2015).

Finally, neuromasts in Honey Creek Cave fish are more numerous than in surface fish, concordant with observations in Mexican cave and surface fish where expansion of neuromasts provides critical spatial and environmental information in dark environments (Yoshizawa et al., 2010). In sum, we observe many shifts in behavior and several changes in morphology between surface fish and the recently established cavefish population that parallel those observed in the long-established Mexican tetras.

In contrast, shifts in several traits in Honey Creek Cave fish were notably discordant with those observed in Mexican cavefish populations (Table 1). First, we found that eye diameter was larger in Honey Creek Cave individuals relative to surface fish when standardized by length of the fish. Fish in our study were all collected within 100 meters of the cave entrance, as fish are not abundant farther into the cave and primarily occur near the twilight zone of cave entrances. While our observation is contrary to most other studies in cavefish (Borowsky, 2015), larger eye size in Honey Creek Cave fish may reflect selection for improved vision in low light conditions, as documented in nocturnal fish (Schmitz and Wainwright, 2011) and organisms that live in the twilight zone (Camp and Jensen, 2007; but see Iglesias et al., 2018). Eye size per unit length was consistent between 2017 and 2018 sampling of Honey Creek Cave fish. However, standardized eye size in Honey Creek Cave fish was larger than in museum collections from Hays County (1976) but similar in size to Kerr County (1979), suggesting that the eye phenotype of the Honey Creek Cave population may require closer morphological inspection (e.g. histology) to detect consistent differences that may exist between cave and surface populations.

Second, in a mirror-elicited aggression assay, Honey Creek Cave fish adhered to the mirror (typically a proxy for aggression) for a higher proportion of time than surface fish, contrary to established Mexican cavefish populations that are less aggressive than surface fish (Burchards et al., 1985; Elipot et al., 2014; Elipot et al., 2013; Espinasa et al., 2005; Hinaux et al., 2015; Langecker et al., 1995; Rétaux and Elipot, 2013). The mirror test is not possible with Mexican cavefish, since they are blind and aggression must be measured by intruder assays (Elipot et al., 2013). We chose the mirror aggression assay rather than an intruder assay to avoid the need to match fish by size and sex. The use of different test types may have influenced the results (Oliveira and Canário, 2011). However, a comparison of the mirror assay and intruder assay in zebrafish revealed high concordance (Way et al., 2015), and increased aggression is seen in low resource caves in other species (Melotto et al., 2019). Part of the increased aggression we observed may simply be response to a novel object, since cavefish demonstrated qualitatively less avoidance of novel objects in VAB trials. Future investigations will assay this by also including a trial with the mirror in the tank, with the opaque side facing the subject. Long-established populations of Mexican cavefish possess neuroanatomical and neurochemical differences suggested to shift cavefish behavior from fighting to feeding including: larger anterior paraventricular hypothalamic nucleus that contains more neurons, higher serotonin, 5-HT, dopamine, and noradrenaline in the forebrain, and about half of the MAO activity (monoamine oxidase) seen in surface fish (Elipot et al., 2014; Elipot et al., 2013). Intriguingly, we see shifts toward aggression and away from feeding in Honey Creek Cave fish, which is the opposite of the pattern observed in Mexican cavefish, and future work should investigate the neuroanatomical and neurochemical differences in Texas fish.

Notably, our work suggests additional phenotypes to examine in future studies. While anecdotal, we observed in the process of anesthetizing fish for neuromast staining that surface fish required ice-bath chilled water to immobilize, whereas cavefish required that water temperature be > 7°C to survive. While this is a qualitative observation, surface fish appear to be able to withstand a cold shock substantially better than cavefish. Stenothermy is common among troglobitic organisms and reduced cold tolerance in Honey Creek Cave fish may represent an additional troglomorphic trait (Barr, 1967; Mermillod-Blondin et al., 2013). Future work will include assaying additional traits known to be increased or enhanced in Mexican cavefish such as number of tastebuds (Varatharasan et al., 2009; Yamamoto et al., 2009), number of teeth (Atukorala et al., 2013; Yamamoto et al., 2003), odorant detection ability (Bibliowicz et al., 2013; Protas et al., 2008), and prey capture skills (Espinasa et al., 2014).

In addition to increasing the number of traits characterized, there is ample opportunity to expand this work to other populations. We have observed *A. mexicanus* in other caves and wells in Central Texas that lack direct connection to surface populations or are only ephemerally connected. A better understanding of the distribution, life history, and ecology of subterranean *A. mexicanus* populations of recent origin may shed light on cave colonization, and is also important from a conservation context. Many stygobitic organisms in the Edwards Aquifer (including federally-listed invertebrates, fish, and salamanders) are threatened by anthropogenic factors, primarily alterations to the quality and quantity of groundwater in the Edwards Aquifer (Page, 2016). The invasion of a new potential predator in *A. mexicanus* could have a pronounced, negative effect on sensitive cave and aquifer ecosystems. This species has been documented at several sites where federally-listed species occur (Gluesenkamp et al., 2018) and has been observed consuming state-listed groundwater species (AGG pers obs). Thus, more intense study of this system would also aid in understanding of the potential threat that *A. mexicanus* pose to native stygobites in Texas.

In conclusion, we have documented a variety of morphological and behavioral trait differences between Honey Creek surface and cave populations of *Astyanax mexicanus* in Central Texas, despite likely recent origin of the cave population. Shifts in several of these traits (e.g. coloration, dorsal fin placement, feeding, and wall-following) are concordant with changes in traits observed in the long-established cave populations in the Sierra de El Abra and Sierra de Guatemala regions in Mexico. Interestingly, we found that some trait shifts are in the opposite direction of those observed in Mexican cavefish populations (e.g., eye size, neuromast size and number, aggression, and body condition). Finally, some traits exhibit no difference between Honey Creek Cave and surface fish (e.g. spatial tank usage in the dark). Notably, we observed a qualitative but striking sensitivity of Honey Creek Cave fish to cold shock. While additional studies of the underlying processes shaping these phenotypes are needed, this population offers a promising and unique opportunity to study the first stages in the colonization of the subterranean environment by a surface organism.

## Data Accessibility Statement

All raw data is available as supplementary files associated with this manuscript.

## Acknowledgements

We thank Joyce Moore for generously providing access to Honey Creek Cave and Texas Parks and Wildlife Department for providing access to Honey Creek State Natural Area. Beverly Burmeier, Aimee Beveridge, Jessica Gordon, Geoff Hoese, Bennett Lee, and Jack, Leah, and Ruby Gluesenkamp provided assistance in the field. We also thank Adam Cohen, Dean Hendrickson (Biodiversity Collections, UT Austin), Kevin Conway, and Toby Hibbitts (Biodiversity and Teaching Collections, Texas A&M) for their generous assistance in accessing UT and TAMU collections and Thomas Pengo, at the University of Minnesota Informatics Institute (UMII) and Mark Sanders at the UMN University Imaging Centers for help imaging neuromasts. Mason Lee, Bekky Muscher-Hodges, and Kamryn Richard provided invaluable logistical support for fieldwork and behavioral trials. Dave Stephens generously donated some of his animal space to house fish. Nate Swanson, Katrina Carrow, Nathaniel Rose, Matthew Chrostek, Ilya Moskalenko, Nicholas Plachinski, and Jeff Miller assisted with fish care, behavioral trials and/or data collection. Rachel Moran and Jeff Miller provided excellent feedback on the manuscript.

## Supplementary Material

### Supplementary Tables

**Table S1.**
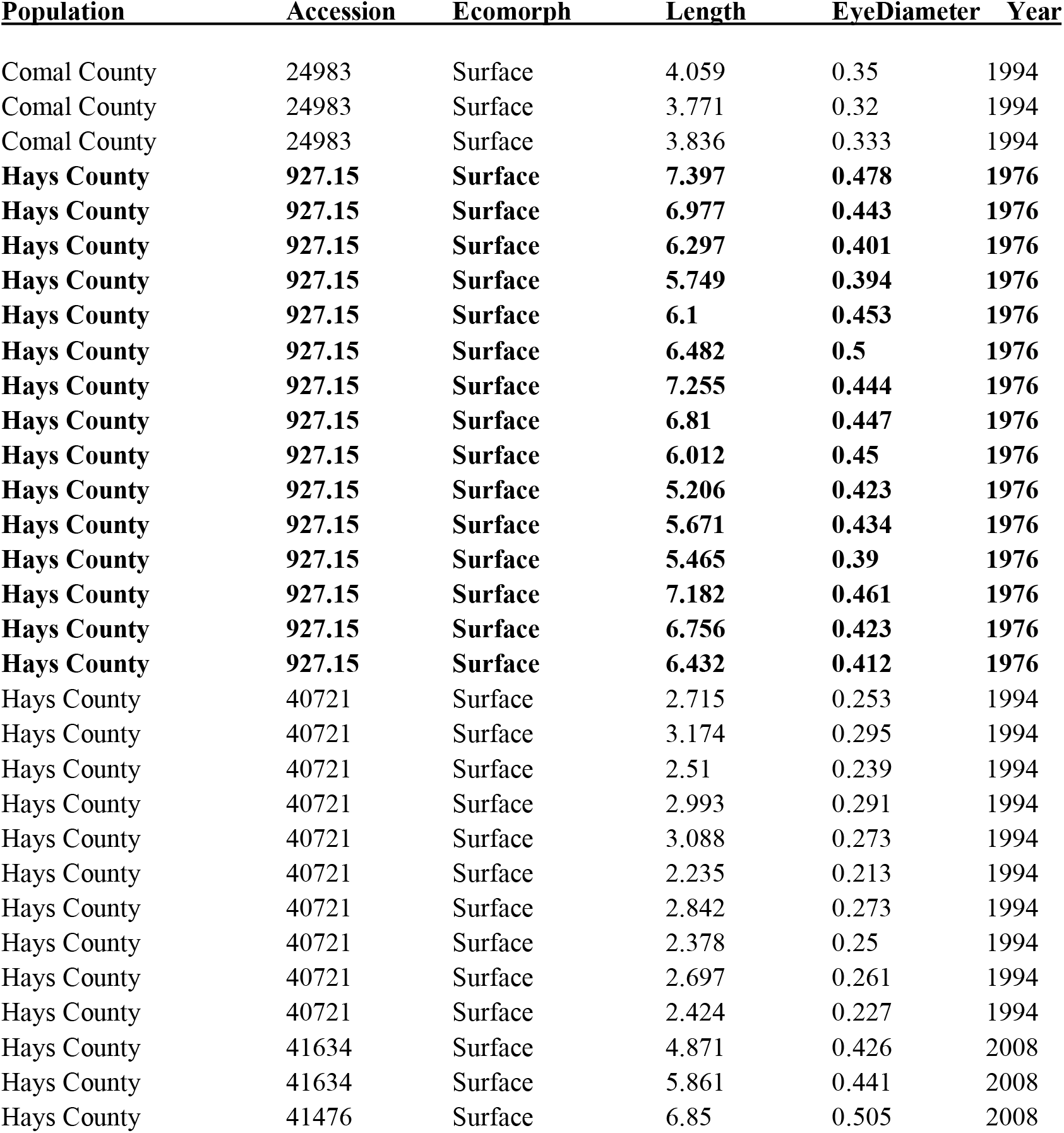
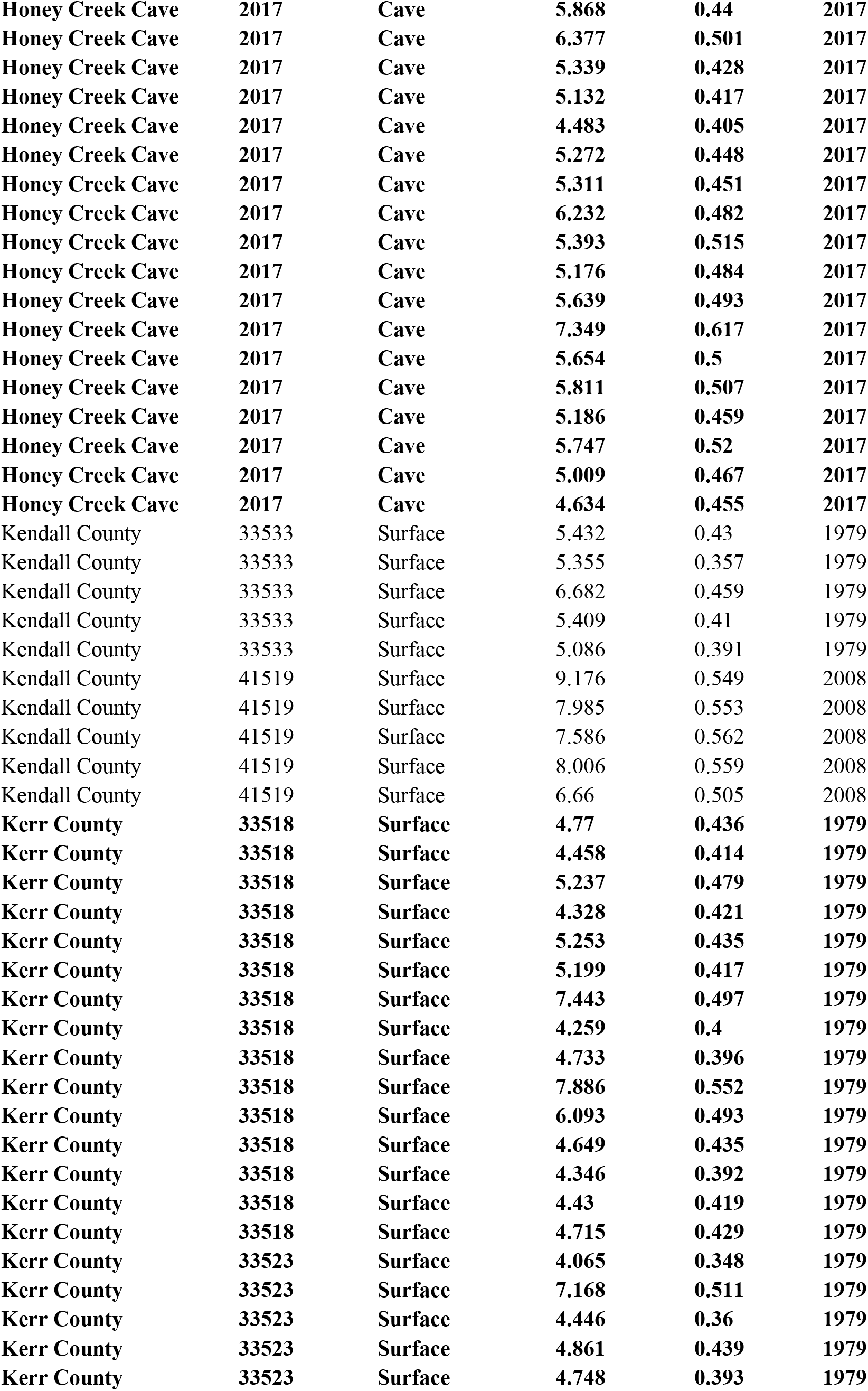
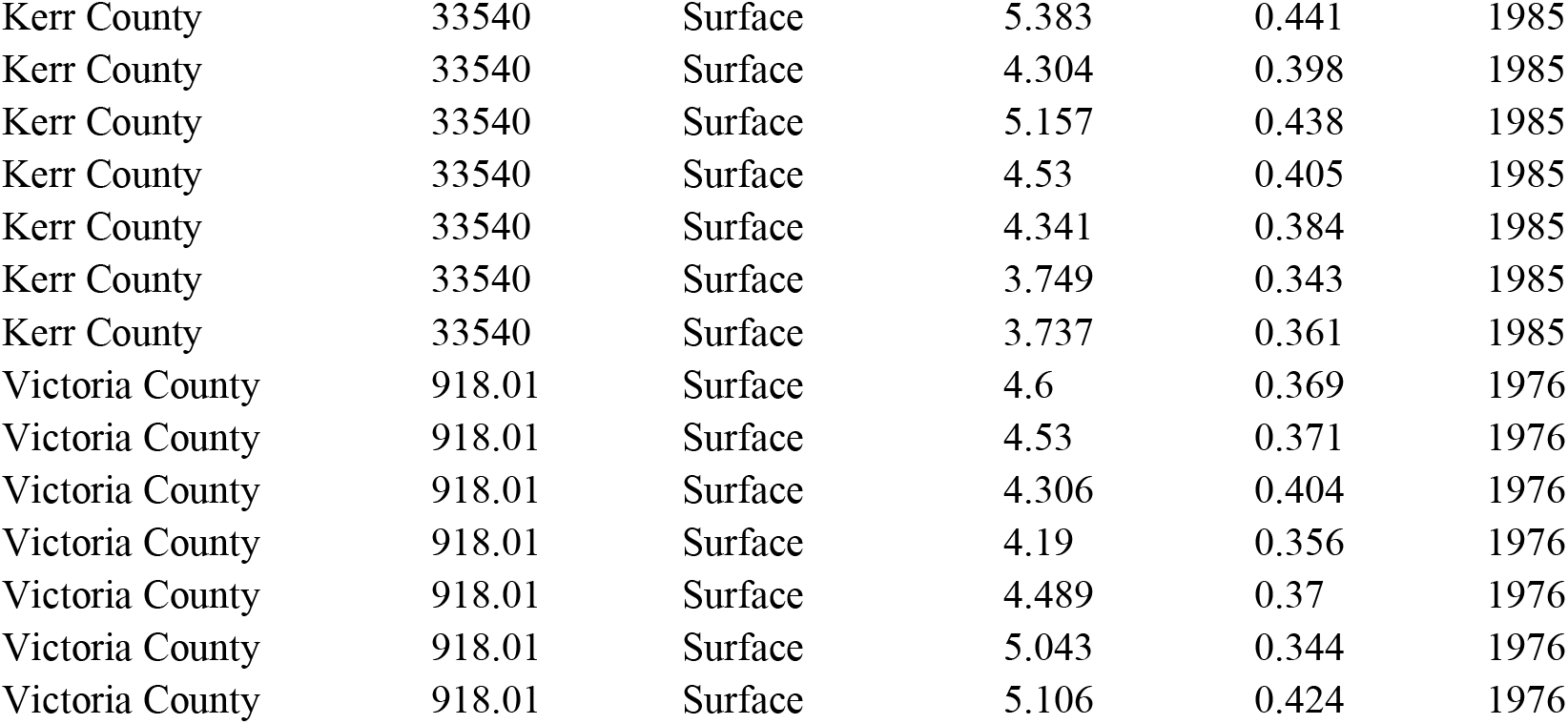
Preserved specimens from the Guadalupe River Basin and date of collection. Samples were measured from University of Texas Austin Texas Memorial Museum and Texas A&M University Biodiversity Research and Teaching Collections. Bold samples were used for analysis in the paper due to small sample sizes for other locality-date combinations and because these samples from Hay Co. and Kerr Co. were taken prior to the first documented *Astyanax* in Honey Creek Cave.

| 70 | **Population** | **Accession** | **Ecomorph** | **Length** | **EyeDiameter** | **Year** |
| --- | --- | --- | --- | --- | --- | --- |
| 71 | Comal County | 24983 | Surface | 4.059 | 0.35 | 1994 |
| 72 | Comal County | 24983 | Surface | 3.771 | 0.32 | 1994 |
| 73 | Comal County | 24983 | Surface | 3.836 | 0.333 | 1994 |
| 74 | **Hays County** | **927.15** | **Surface** | **7.397** | **0.478** | **1976** |
| 75 | **Hays County** | **927.15** | **Surface** | **6.977** | **0.443** | **1976** |
| 76 | **Hays County** | **927.15** | **Surface** | **6.297** | **0.401** | **1976** |
| 77 | **Hays County** | **927.15** | **Surface** | **5.749** | **0.394** | **1976** |
| 78 | **Hays County** | **927.15** | **Surface** | **6.1** | **0.453** | **1976** |
| 79 | **Hays County** | **927.15** | **Surface** | **6.482** | **0.5** | **1976** |
| 80 | **Hays County** | **927.15** | **Surface** | **7.255** | **0.444** | **1976** |
| 81 | **Hays County** | **927.15** | **Surface** | **6.81** | **0.447** | **1976** |
| 82 | **Hays County** | **927.15** | **Surface** | **6.012** | **0.45** | **1976** |
| 83 | **Hays County** | **927.15** | **Surface** | **5.206** | **0.423** | **1976** |
| 84 | **Hays County** | **927.15** | **Surface** | **5.671** | **0.434** | **1976** |
| 85 | **Hays County** | **927.15** | **Surface** | **5.465** | **0.39** | **1976** |
| 86 | **Hays County** | **927.15** | **Surface** | **7.182** | **0.461** | **1976** |
| 87 | **Hays County** | **927.15** | **Surface** | **6.756** | **0.423** | **1976** |
| 88 | **Hays County** | **927.15** | **Surface** | **6.432** | **0.412** | **1976** |
| 89 | Hays County | 40721 | Surface | 2.715 | 0.253 | 1994 |
| 90 | Hays County | 40721 | Surface | 3.174 | 0.295 | 1994 |
| 91 | Hays County | 40721 | Surface | 2.51 | 0.239 | 1994 |
| 92 | Hays County | 40721 | Surface | 2.993 | 0.291 | 1994 |
| 93 | Hays County | 40721 | Surface | 3.088 | 0.273 | 1994 |
| 94 | Hays County | 40721 | Surface | 2.235 | 0.213 | 1994 |
| 95 | Hays County | 40721 | Surface | 2.842 | 0.273 | 1994 |
| 96 | Hays County | 40721 | Surface | 2.378 | 0.25 | 1994 |
| 97 | Hays County | 40721 | Surface | 2.697 | 0.261 | 1994 |
| 98 | Hays County | 40721 | Surface | 2.424 | 0.227 | 1994 |
| 99 | Hays County | 41634 | Surface | 4.871 | 0.426 | 2008 |
| 100 | Hays County | 41634 | Surface | 5.861 | 0.441 | 2008 |
| 101 | Hays County | 41476 | Surface | 6.85 | 0.505 | 2008 |
| 102 | Honey Creek Cave | 2017 | Cave | 5.868 | 0.44 | 2017 |
| 103 | Honey Creek Cave | 2017 | Cave | 6.377 | 0.501 | 2017 |
| 104 | Honey Creek Cave | 2017 | Cave | 5.339 | 0.428 | 2017 |
| 105 | Honey Creek Cave | 2017 | Cave | 5.132 | 0.417 | 2017 |
| 106 | Honey Creek Cave | 2017 | Cave | 4.483 | 0.405 | 2017 |
| 107 | Honey Creek Cave | 2017 | Cave | 5.272 | 0.448 | 2017 |
| 108 | Honey Creek Cave | 2017 | Cave | 5.311 | 0.451 | 2017 |
| 109 | Honey Creek Cave | 2017 | Cave | 6.232 | 0.482 | 2017 |
| 110 | Honey Creek Cave | 2017 | Cave | 5.393 | 0.515 | 2017 |
| 111 | Honey Creek Cave | 2017 | Cave | 5.176 | 0.484 | 2017 |
| 112 | Honey Creek Cave | 2017 | Cave | 5.639 | 0.493 | 2017 |
| 113 | Honey Creek Cave | 2017 | Cave | 7.349 | 0.617 | 2017 |
| 114 | Honey Creek Cave | 2017 | Cave | 5.654 | 0.5 | 2017 |
| 115 | Honey Creek Cave | 2017 | Cave | 5.811 | 0.507 | 2017 |
| 116 | Honey Creek Cave | 2017 | Cave | 5.186 | 0.459 | 2017 |
| 117 | Honey Creek Cave | 2017 | Cave | 5.747 | 0.52 | 2017 |
| 118 | Honey Creek Cave | 2017 | Cave | 5.009 | 0.467 | 2017 |
| 119 | Honey Creek Cave | 2017 | Cave | 4.634 | 0.455 | 2017 |
| 120 | Kendall County | 33533 | Surface | 5.432 | 0.43 | 1979 |
| 121 | Kendall County | 33533 | Surface | 5.355 | 0.357 | 1979 |
| 122 | Kendall County | 33533 | Surface | 6.682 | 0.459 | 1979 |
| 123 | Kendall County | 33533 | Surface | 5.409 | 0.41 | 1979 |
| 124 | Kendall County | 33533 | Surface | 5.086 | 0.391 | 1979 |
| 125 | Kendall County | 41519 | Surface | 9.176 | 0.549 | 2008 |
| 126 | Kendall County | 41519 | Surface | 7.985 | 0.553 | 2008 |
| 127 | Kendall County | 41519 | Surface | 7.586 | 0.562 | 2008 |
| 128 | Kendall County | 41519 | Surface | 8.006 | 0.559 | 2008 |
| 129 | Kendall County | 41519 | Surface | 6.66 | 0.505 | 2008 |
| 130 | Kerr County | 33518 | Surface | 4.77 | 0.436 | 1979 |
| 131 | Kerr County | 33518 | Surface | 4.458 | 0.414 | 1979 |
| 132 | Kerr County | 33518 | Surface | 5.237 | 0.479 | 1979 |
| 133 | Kerr County | 33518 | Surface | 4.328 | 0.421 | 1979 |
| 134 | Kerr County | 33518 | Surface | 5.253 | 0.435 | 1979 |
| 135 | Kerr County | 33518 | Surface | 5.199 | 0.417 | 1979 |
| 136 | Kerr County | 33518 | Surface | 7.443 | 0.497 | 1979 |
| 137 | Kerr County | 33518 | Surface | 4.259 | 0.4 | 1979 |
| 138 | Kerr County | 33518 | Surface | 4.733 | 0.396 | 1979 |
| 139 | Kerr County | 33518 | Surface | 7.886 | 0.552 | 1979 |
| 140 | Kerr County | 33518 | Surface | 6.093 | 0.493 | 1979 |
| 141 | Kerr County | 33518 | Surface | 4.649 | 0.435 | 1979 |
| 142 | Kerr County | 33518 | Surface | 4.346 | 0.392 | 1979 |
| 143 | Kerr County | 33518 | Surface | 4.43 | 0.419 | 1979 |
| 144 | Kerr County | 33518 | Surface | 4.715 | 0.429 | 1979 |
| 145 | Kerr County | 33523 | Surface | 4.065 | 0.348 | 1979 |
| 146 | Kerr County | 33523 | Surface | 7.168 | 0.511 | 1979 |
| 147 | Kerr County | 33523 | Surface | 4.446 | 0.36 | 1979 |
| 148 | Kerr County | 33523 | Surface | 4.861 | 0.439 | 1979 |
| 149 | Kerr County | 33523 | Surface | 4.748 | 0.393 | 1979 |
| 150 | Kerr County | 33540 | Surface | 5.383 | 0.441 | 1985 |
| 151 | Kerr County | 33540 | Surface | 4.304 | 0.398 | 1985 |
| 152 | Kerr County | 33540 | Surface | 5.157 | 0.438 | 1985 |
| 153 | Kerr County | 33540 | Surface | 4.53 | 0.405 | 1985 |
| 154 | Kerr County | 33540 | Surface | 4.341 | 0.384 | 1985 |
| 155 | Kerr County | 33540 | Surface | 3.749 | 0.343 | 1985 |
| 156 | Kerr County | 33540 | Surface | 3.737 | 0.361 | 1985 |
| 157 | Victoria County | 918.01 | Surface | 4.6 | 0.369 | 1976 |
| 158 | Victoria County | 918.01 | Surface | 4.53 | 0.371 | 1976 |
| 159 | Victoria County | 918.01 | Surface | 4.306 | 0.404 | 1976 |
| 160 | Victoria County | 918.01 | Surface | 4.19 | 0.356 | 1976 |
| 161 | Victoria County | 918.01 | Surface | 4.489 | 0.37 | 1976 |
| 162 | Victoria County | 918.01 | Surface | 5.043 | 0.344 | 1976 |
| 163 | Victoria County | 918.01 | Surface | 5.106 | 0.424 | 1976 |
| 164 |  |  |  |  |  |  |

**Table S2.** Wilcoxon test W and p-values for all tests for live fish to determine which color components are different between cave and surface fish. Significant values are in bold.

|  | Anal | Dorsal | Middle | Tail |
| --- | --- | --- | --- | --- |
| R | 375.5, p = 0.684 | **502.5, p = 0.009** | 397, p = 0.436 | 403, p = 0.377 |
| G | 378, p = 0.653 | 431, p = 0.172 | 419, p = 0.249 | 418, p = 0.254 |
| B | 350, p = 0.986 | 346, p = 0.931 | 363, p = 0.849 | 396.5, p = 0.441 |
| H | 255.5, p = 0.098 | 250, p = 0.080 | 272, p = 0.172 | 245, p = 0.066 |
| S | **629.5, p < 0.001** | **665.5, p < 0.001** | **607, p < 0.001** | **577, p < 0.001** |
| L | 384, p = 0.574 | 437, p = 0.141 | 397.5, p = 0.431 | 393.5, p = 0.473 |

**Table S3.** Wilcoxon test and p-values for all tests for dead fish to determine which color components are different between cave and surface fish. Significant values are in bold.

|  | Anal | Dorsal | Middle | Tail |
| --- | --- | --- | --- | --- |
| R | **171, p = 0.032** | 153, p = 0.155 | **27, p < 0.001** | **192, p < 0.001** |
| G | 165, p = 0.052 | 137, p = 0.435 | **24, p < 0.001** | **181, p = 0.011** |
| B | 133, p = 0.535 | **50, p = 0.008** | **22, p < 0.001** | 115.5, p = 0.968 |
| H | **216, p < 0.001** | **198, p = 0.001** | 79, p = 0.133 | 179, p = 0.014 |
| S | **231, p < 0.001** | **227, p < 0.001** | **214.5, p < 0.001** | **233, p < 0.001** |
| L | **169, p = 0.039** | 137.5, p = 0.423 | **26, p < 0.001** | **183, p = 0.009** |

**Table S4.** Numbers are mean with median in parentheses. Paired one-tailed Wilcoxon Tests for VAB trials.

190 **Number of Approaches**
|  | **Trial 1** | **Trial 2** | **Trial 3** |
| --- | --- | --- | --- |
| 192 **Cave** | NA | 4.72 (5.00) | 5.00 (4.50) |
| 193 **Surface** | NA | 4.00 (3.00) | 4.28 (4.00) |

|  | **Trial 1** | **Trial 2** | **Trial 3** |
| --- | --- | --- | --- |
| 200 **Cave** | 5.74% (5.67%) | 6.58% (5.05%) | 11.17% (3.50%) |
| 201 **Surface** | 18.80% (16.18%) | 12.66% (8.15%) | 14.87% (11.64%) |

|  | **Trial 1** | **Trial 2** | **Trial 3** |
| --- | --- | --- | --- |
| 210 **Cave** | 6.21 (6.50) | 7.21 (7.50) | 6.00 (4.50) |
| 211 **Surface** | 10.59 (11.00) | 7.41 (7.00) | 8.05 (7.50) |

Cave Trial 1 - Trial 2 V = 38, p-value = 0.713
Cave Trial 1 - Trial 3 V = 46.5, p-value = 0.290
\*Surface Trial 1 - Trial 2 V = 470.5, p-value < 0.001
\*Surface Trial 1 - Trial 3 V = 330, p-value = 0.053

**Table S5.** Mean and medians (in parentheses) for values tracked by Ethovision. For dark trials and light trials, no variables exhibited a significant difference between cave and surface fish. However, in a paired analysis both cave and surface fish spent more time at the bottom of the tank in the light trials. Surface fish significantly slowed their velocity and reduced their distance traveled. We analyzed 15 cavefish and 37 surface fish. Asterisks represent values that were different bewteen light and dark trials.

**Dark Trial**
|  | Proportion |  |  | Proportion time on |
| --- | --- | --- | --- | --- |
|  | time immobile | distance traveled(cm) | Velocity (cm/s) | bottom |
| Cave | 0.14 (0.09) | 1667.90 (1667.84) | 5.67 (5.66) | 0.51 (0.45) |
| Surface | 0.20 (0.06) | 1841.69 (1733.84) | 6.20 (5.83) | 0.55 (0.56) |

**Lighted Trial**
|  | Proportion |  |  | Proportion time on |
| --- | --- | --- | --- | --- |
|  | time immobile | distance traveled(cm) | Velocity (cm/s) | bottom |
| Cave | 0.18 (0.09) | 1639.92 (1700.88) | 5.790 (6.36) | 0.80 (0.84)* |
| Surface | 0.25 (0.18) | 1556.46 (1581.81)* | 5.307 (5.91)* | 0.77 (0.79)* |

### Supplementary Figures

**Figure S1.**
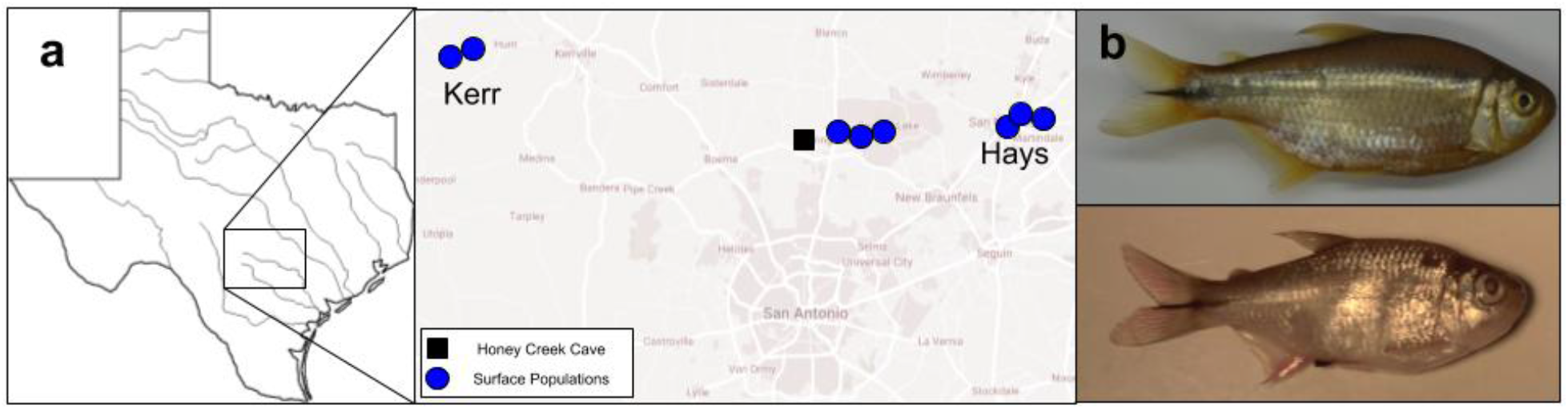
Sampling sites of all cave and surface individuals included in Table S1.

**Figure S2.**
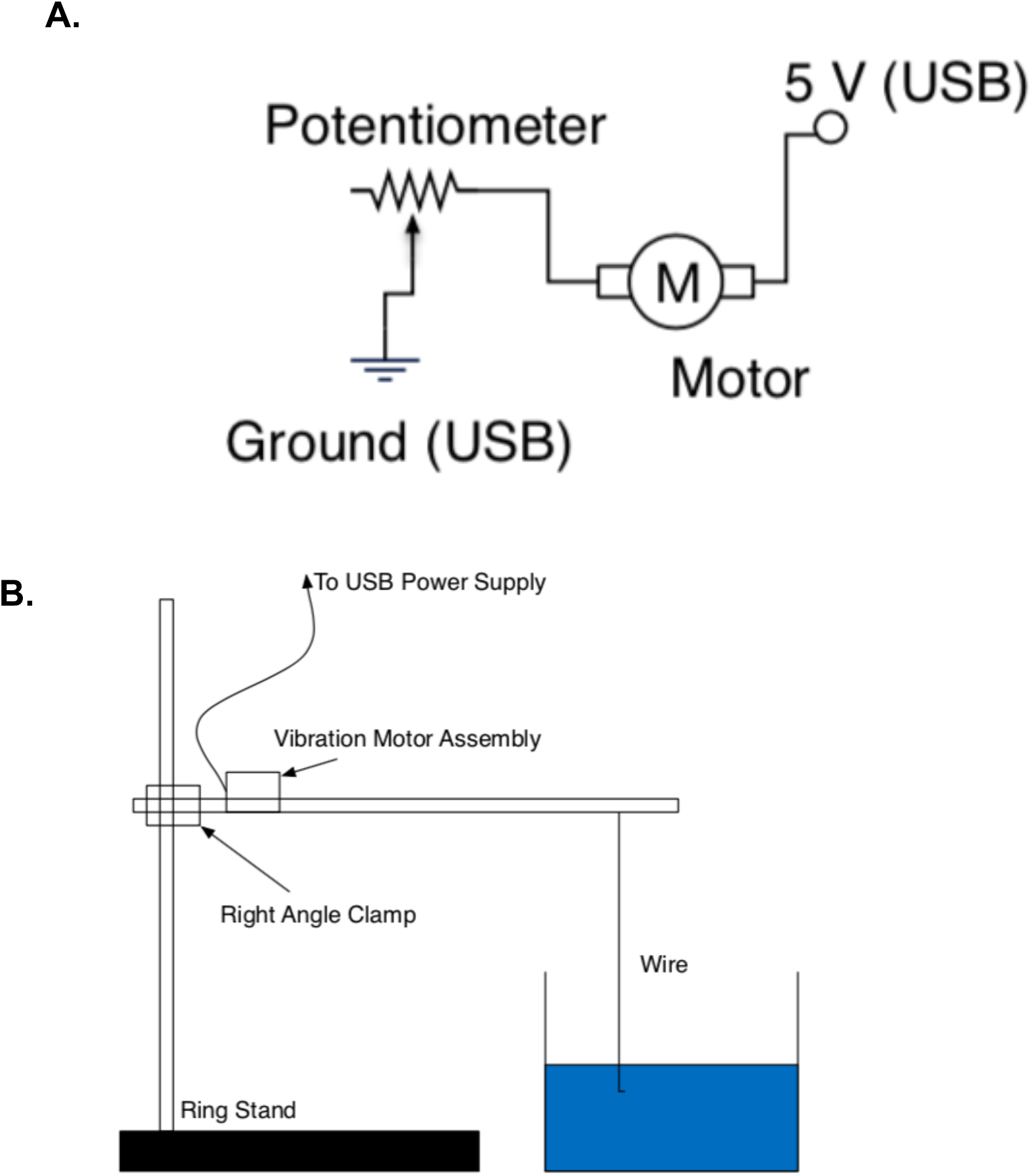
A. Circuit diagram of the excitation mechanism connected to the plastic rods. **B.** Set-up for the VAB trials.

